# Single Cell Transcriptomics and Surface Protein Expression Reveal Distinct Cellular and Molecular Phenotypes in Human RPESC-RPE and PSC-RPE

**DOI:** 10.64898/2026.02.26.708319

**Authors:** Swapna Nandakumar, Farhad Farjood, Taylor Bertucci, Steven Lotz, Skanda Sai, Yue Wang, Jade A. Kozak, Brigitte L. Arduini, Jeffrey H. Stern, Nathan C. Boles, Sally Temple

## Abstract

Current retinal pigment epithelium (RPE) cell replacement strategies in trials for age-related macular degeneration (AMD) are based on either pluripotent stem cell (PSC) or adult RPE stem cell (RPESC) sources. We used Cellular Indexing of Transcriptomes and Epitopes by Sequencing (CITE-Seq) to simultaneously assess single-cell transcriptomic and surface protein information, comparing these two RPE sources. Both RPESC-RPE and PSC-RPE expressed key RPE markers and exhibited cellular heterogeneity. However, RPESC-RPE had higher expression of genes related to mature retinal functions, whereas PSC-RPE had greater expression of genes involved in stem cell development and differentiation. We identified two surface proteins that distinguished the cell types. The “don’t eat me” signal, CD24, was detected robustly on adult RPESC-RPE cells, while CD57 was detected on most PSC-RPE cells. The differences in gene and surface protein expression suggest that the two RPE sources differ in functional, adhesion, and immunomodulatory properties, which may impact transplantation outcomes.

## INTRODUCTION

The retinal pigment epithelium (RPE) is the outermost layer of the retina and plays a crucial role in visual function. It is a monolayer of polarized, pigmented epithelial cells situated between Bruch’s membrane and the choroidal vasculature basally and photoreceptor outer segments apically. The RPE functions as a blood-retina barrier and transports nutrients and small molecules required for photoreceptor function.^1^ It plays a key role in the visual cycle by converting all-trans-retinol to 11-cis-retinol, which is necessary for the activation of opsins in photoreceptors.^2^ The RPE also phagocytoses and renews the photoreceptor outer segments. A healthy RPE is essential for maintaining the health of photoreceptors and visual function.^3^ Significant RPE dysfunction and drusen deposits are hallmarks of dry AMD. As AMD progresses, loss of RPE occurs, predominantly in the macula, the central retina responsible for high acuity and color vision. This causes photoreceptor dysfunction and visual impairment. Dry AMD can progress to geographic atrophy, characterized by the loss of both photoreceptors and the underlying choroid, leading to severe loss of central vision.^4^ In approximately 10–20% of AMD cases, neovascularization occurs due to the invasion of the choriocapillaris into the RPE, resulting in wet AMD (also termed neovascular or exudative AMD).^5^ Anti-VEGF therapy helps with the management of wet AMD.^6^ However, there are presently no vision-improving treatment options available for early-stage dry AMD.

One therapeutic strategy to improve vision involves restoring the RPE layer by transplanting new RPE cells to augment or replace the damaged epithelium. Based on information available from trial registries, there are 83 pluripotent stem cell (PSC)-derived products in clinical trials, with 17 of these targeting RPE replacement, across the USA, Asia, and Europe.^7^ PSC-RPE and adult stem cell–derived RPESC-RPE are the two primary RPE sources in clinical trials. However, similarities and differences between the RPE generated from PSC and adult stem cell sources remain to be defined, motivating this comparative study.

Several ongoing clinical trials utilize an RPE cell suspension; others use a monolayer patch of RPE cells on a scaffold.^8^ In the case of cells on a scaffold, the goal is to place the RPE cells subretinally in a position to interact with the photoreceptor layer, while adapting to the curvature of the retina and minimally altering the focal length of the eye with the added scaffold/cell layer.^9,10^ When a cell suspension is used, the goal is for the cells to integrate into the existing host RPE layer and to augment RPE functions. Both approaches have been achieved in animal models.^10–15^ Successful cell transplantation using either approach depends on the ability of donor cells to engraft into the retina, integrate with host tissue, and restore function.^16^ Cell membrane proteins play a key role in graft success. They mediate cell-cell and cell-environment interactions, playing a key role in cell survival, adhesion, immune events, phagocytosis, growth, and proliferation.^17^ Surface markers can also be used to distinguish cell subtypes and enrich desired cell populations to improve transplantation success.^18,19^ Given their diversity of function and potential therapeutic impact, identifying surface proteins is important.

In this study, we cultured PSC-RPE and adult RPESC-RPE under identical conditions, and applied CITE-Seq,^20^ which integrates single-cell transcriptomics with surface protein profiling to achieve an in-depth comparative characterization. Our goal was to identify pathways and surface molecules that could inform strategies to improve clinical outcomes. Both RPE sources expressed canonical RPE markers, consistent with prior reports.^21,22^ However, our analysis revealed significant differences in the single-cell transcriptome and surfaceome, suggesting intrinsic biologic distinctions that may contribute to different outcomes post-transplantation.

## RESULTS

### Single-cell RNA sequencing reveals similar transcriptomic clusters in adult RPESC-RPE and PSC-RPE

RPESC-RPE cells were derived from three eye donor and maintained in culture as described previously (**Supplementary Figure 1a, Table S1**).^21,23^ PSC-RPE cells were derived from three different donor human PSC (hPSC) lines using a previously published protocol (**Supplementary Figure 1b, Table S1**).^24^ Prior studies demonstrated that RPESC-RPE and PSC-RPE generated with these two methods exhibit canonical RPE markers and functions, including the formation of tight junctions, phagocytosis of photoreceptor outer segments, and polarized release of PEDF and VEGF.^21,22^ All experiments were conducted using RPESC-RPE or PSC-RPE cells that were thawed at passage 2 and then cultured for an additional 10 weeks under the same RPE maintenance media and culture conditions. The resulting cultures had an overall similar appearance, albeit with RPESC-RPE being slightly larger than PSC-RPE (**Supplementary Figure 1c, Table S2**). Despite the difference in cell sizes, both RPESC-RPE and PSC-RPE exhibited similar aspect ratios (AR), which define whether a cell is regular (AR=1) or has uneven sides (AR=3),^25^ and had similar numbers of neighbors (**Supplementary Figure 1d,e, Table S2**). The RPE cells from each source had an average AR of 1.5 and an average number of neighbors between 5-7, typical of a monolayer of hexagonal-shaped cells.

We examined differences at single-cell resolution between the two cell types using CITE-seq, a sequencing pipeline that multiplexes cell surface protein detection using DNA-tagged antibodies with transcriptomic profiling (**Figure 1a**). A panel of 164 barcoded antibodies, including 4 isotype controls (**Table S3**), was used to track surface protein expression. A total of 2,730 PSC-RPE and 2,504 RPESC-RPE cells were used for the single cell analyses. The median depth of the transcriptomic reads was ∼76,000 reads per cell, detecting a median of ∼2,700 features. The median read depth of the antibody-derived tags (ADT) was ∼32,700 counts per cell. The data from each cell source was integrated via the reciprocal principal component analysis (RPCA) approach in the Seurat package.

**Figure 1:**
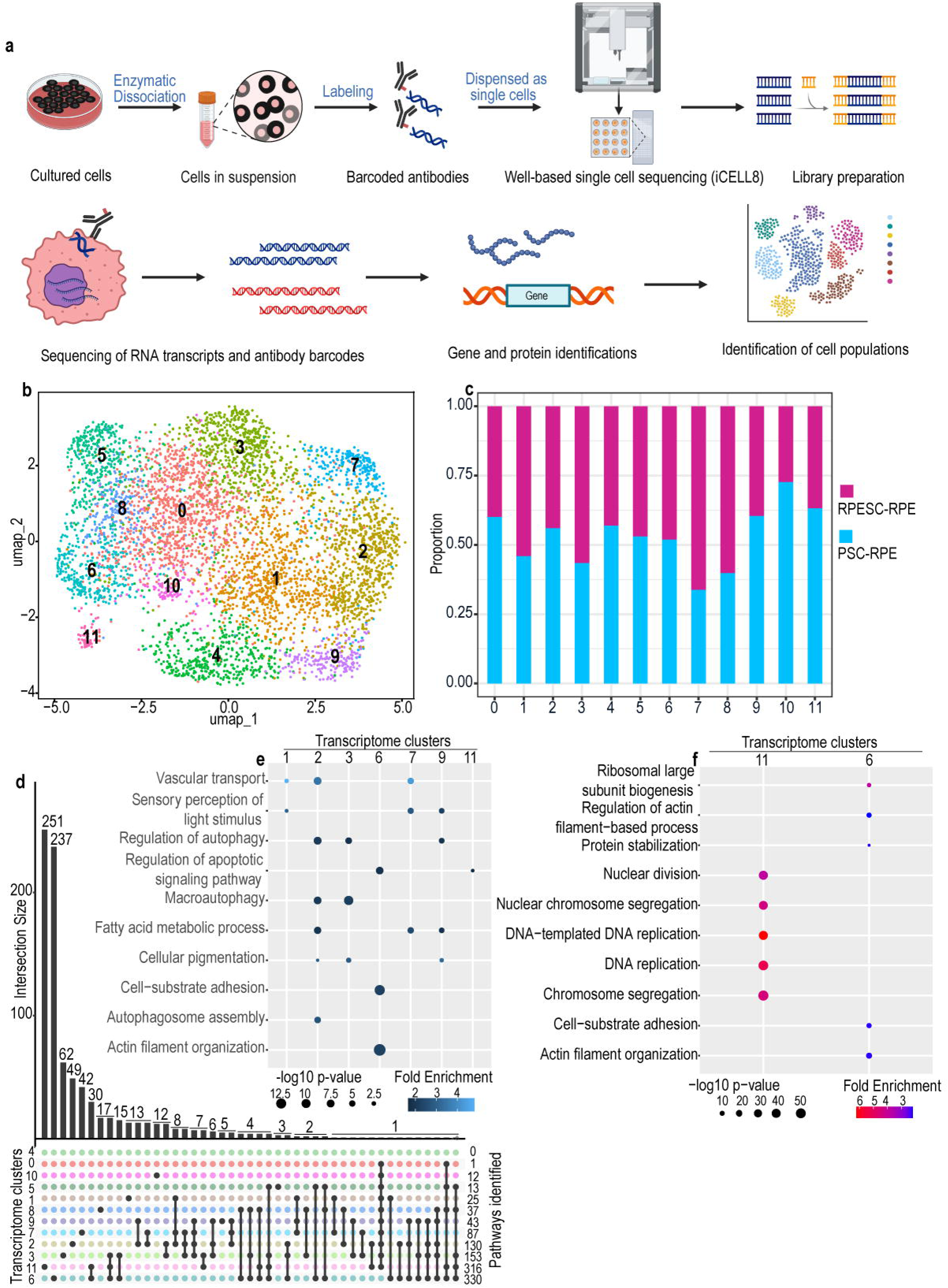
Single cell clustering of RPESC-RPE and PSC-RPE based on transcriptome data. (a) CITE-seq workflow. (b) DimPlot showing clustering of RPESC-RPE and PSC-RPE cells based on single cell transcriptome profile, clusters identified by numbers 0-11. (c) Proportion of RPESC-RPE and PSC-RPE in each transcriptome cluster. (d) Upset plot showing the distribution in overlap of enriched pathways across transcriptome clusters. (e) DotPlot showing pathway enrichment for GO-BP terms related to RPE functions across transcriptome clusters. (f) DotPlot showing top 5 unique enriched pathways for transcriptome clusters 11 and 6.

First, we focused on the single-cell transcriptome data. We identified 12 clusters in our transcriptomic dataset. A 2-dimensional uniform manifold approximation and projection (UMAP) of the integrated RNA-seq data that included both RPESC-RPE and PSC-RPE cells did not demonstrate a clear separation between the two cell sources, with both cell sources contributing to each identified cluster (**Figure 1b,c**). To investigate potential regional identity, we examined the expression of genes previously reported to be differentially expressed in macular versus temporal/nasal regions of RPE/choroid preparations. Cells across all clusters expressed both macular and peripheral markers (**Table S4**).^26,27^

A prior study established a set of 154 RPE signature genes based on bulk analysis of native fetal and adult RPE that are linked to key RPE functions, including sensory and light perception, pigment biosynthesis, phagocytic activity, and transporter activity.^28^ Of these signature genes,149 were detected in all clusters, and clusters 0, 1, 2, 3, and 5 expressed all 154 (**Supplementary Figure 1f, Table S5**). Hence, both adult RPESC- and PSC-derived cultures were confirmed to be highly enriched for bona fide RPE cells, exhibiting comparable transcriptional profiles that clustered similarly. Among the RPE signature genes, *DHPS,* which is involved in the hypusination of the translation initiation factor eIF5A,^29^ was not detected in clusters 4, 6, 7, 8, and 9 (**Supplementary Figure 1f, Table S5**). In cluster 10, both *DHPS* and *SLC6A15* were undetected, while cluster 11 lacked expression of five genes: *DHPS, FOXD1, PLAG1, SLC4A2,* and *SLC6A15* (**Supplementary Figure 1f, Table S5**).

A single-cell transcriptomic analysis of PSC-RPE cells conducted by Parikh et al.,^22^ identified an RPE subpopulation defined by expression of the transcription factors *FOS*, *JUND*, and *MAFF*, which emerged only after transplantation and was associated with a “terminally mature” RPE state. These genes were proposed to regulate networks that enhance RPE survival and promote photoreceptor health after transplantation.^22^ In our dataset, *FOS* and *JUND* were detected in both RPESC- and PSC-derived RPE cells across most of the 12 clusters (**Supplementary Figure 1g**). Specifically, *FOS* was expressed in 45% of RPESC-RPE cells and 32% of PSC-RPE cells, while *JUND* was expressed in 30% and 27% of these cells, respectively.

Taken together, these findings demonstrate that both PSC-RPE and adult RPESC-RPE expressed core RPE signature genes, generated both macular and peripheral RPE without a strong bias towards either identity, and exhibited transcriptional features previously associated with favorable post-transplantation outcomes.

### RPE clusters indicate functional specializations independent of cell source

Next, we performed a more detailed analysis of the RPE clusters to identify differences in biological pathways based on Gene Ontology (GO) biological processes (GO-BP) terms. Differentially expressed genes (DEGs) were identified using the Seurat FindAllMarkers function (**Table S6**). The resulting gene lists were used for pathway enrichment analysis using ClusterProfiler.^30^ This revealed distinct GO-BP enrichments across clusters (**Table S7**). The DEGs mapped to over 300 GO-BP terms across all clusters. Clusters 6 and 11 exhibited the highest number of unique enriched pathways, while no significant pathway enrichment was detected in cluster 4 (**Figure 1d**). Focusing on GO-BP terms that are relevant to canonical RPE functions, we observed evidence of functional specialization among clusters (**Figure 1e**). For example, clusters 2, 3, and 9 were enriched for pathways related to autophagy and cellular pigmentation; clusters 1, 2, and 7 showed enrichment for vascular transport processes; cluster 1 was enriched for transport and non-retinal cell development; clusters 2 and 9 for lipid metabolism; cluster 3 for mitochondrial function and energy generation; cluster 5 for RNA processing and sensory system development; cluster 7 for transport and metabolism; cluster 8 for hormone response and hormone metabolism; and cluster 10 for ion transport. Of particular interest were clusters 11 and 6 due to their high number of unique pathways (**Figure 1d, Table S7**). Cluster 6, which contained the second highest number of unique pathway terms (**Figure 1d**), was evenly composed of RPESC-RPE and PSC-RPE cells (**Figure 1c**) and was uniquely enriched for GO terms related to cell-substrate adhesion and actin filament organization (**Figure 1f, Table S7**). Adhesion-related terms were absent from all other clusters, suggesting both RPE sources contain a specialized subpopulation with enhanced adhesive capacity. This will be interesting to pursue further, as such a subpopulation may better support cell engraftment and barrier formation.^31^ Cluster 11, with the highest number of unique pathway terms, was composed of ∼60% PSC-RPE and ∼40% RPESC-RPE (**Figure 1c**). It was uniquely enriched for cell division and DNA replication pathways (**Figure 1f, Table S7**), which were not detected in other clusters. These features suggest it may include a progenitor-like subpopulation. This is consistent with prior evidence of an progenitor population in the human eye that persists to old age and is capable of being activated *in vitro* into a self-renewing RPESC.^32^ Interestingly, gene sets related to the regulation of apoptosis were enriched only in these two clusters, potentially reflecting the influence of adhesive and proliferative capacity on cell survival. In summary, these transcriptomic comparisons demonstrate cellular and molecular heterogeneity in both RPESC-RPE and PSC-RPE, and suggest that functional states, rather than cell source, are the primary drivers of clustering at the single-cell transcriptomic level.

### Multimodal CITE-seq analysis enhances the separation of RPESC-RPE and PSC-RPE clusters

To better define subpopulation composition in RPESC-RPE and PSC-RPE cells, we performed multimodal clustering, which combines transcriptomic and ADT data, to detect surface protein expression (**Figure 1a**, bottom panel). We applied the weighted nearest neighbor (WNN) analysis, which constructs modality-specific neighbor graphs, assigns modality weights, and integrates them to cluster cells based on both RNA and protein features.^33^ WNN analysis identified 22 multimodal clusters (MO–M21, **Figure 2a**), now showing improved separation between RPESC-RPE and PSC-RPE cells (**Figure 2b**). This indicates that surface protein detection contributes more strongly than transcriptomic data to distinguishing cell origin. To further resolve lineage-specific clusters, we applied DA-seq^34^ and identified 26 origin-enriched subclusters: 12 for PSC-RPE (P1–P12) and 14 for RPESC-RPE (A13–A26), with cell counts ranging from 18 to 315 (**Figure 2c**). Grey dots represent cells within parent clusters that are not enriched for a specific origin in their local neighborhood.

**Figure 2:**
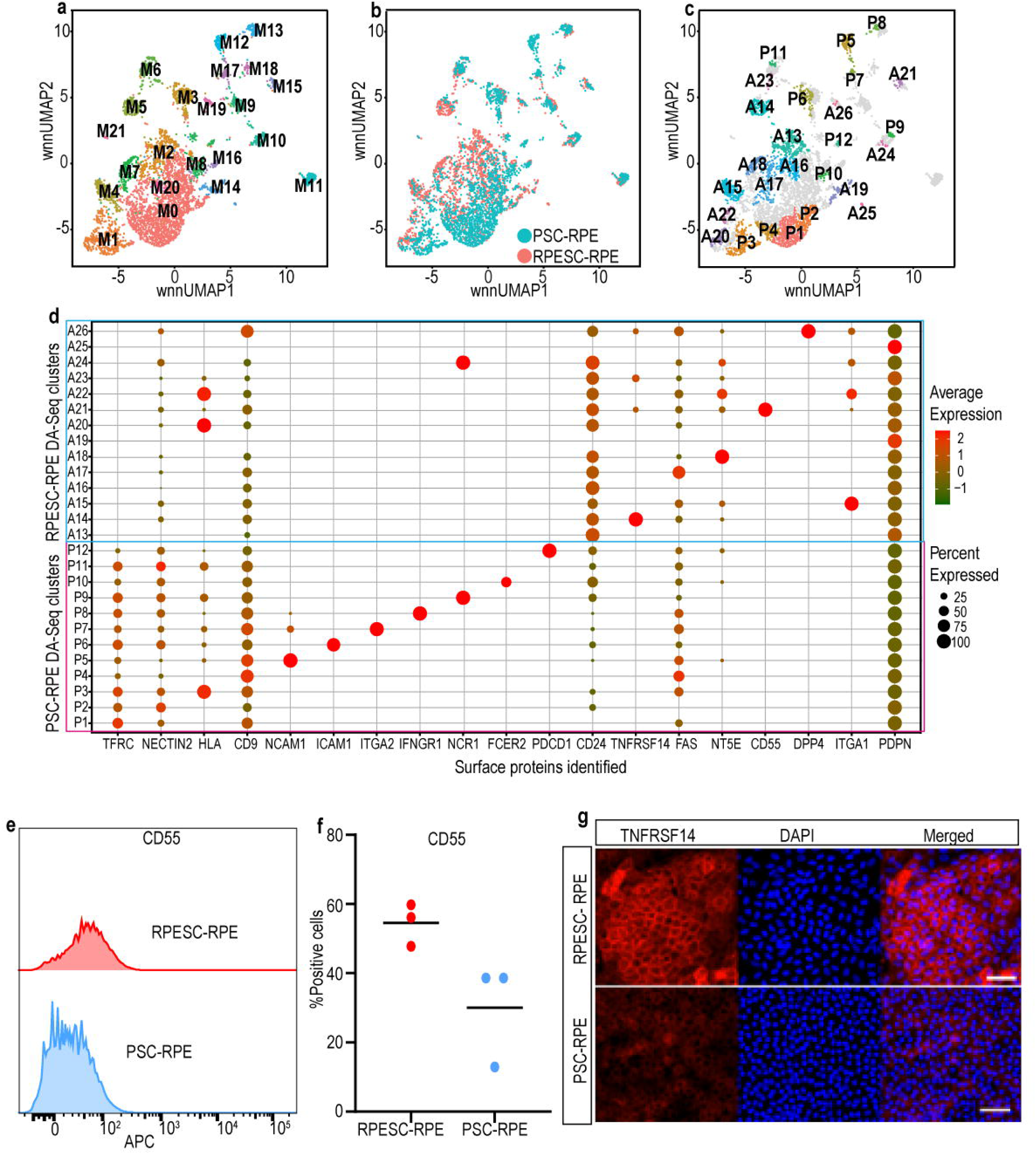
Single cell clustering of RPESC-RPE and PSC-RPE based on multimodal analysis incorporating transcript and surface protein expression. (a) DimPlot of multimodal clustering of adult and PSC-RPE cells, clusters identified by labels M0-M21. (b) DimPlot displaying cell source for multimodal clusters. (c) DimPlot of DA-seq clustering based on cell source origin. Clusters P1-P12 are PSC-RPE biased while clusters A13-A26 are RPESC-RPE biased. (d) DotPlot showing expression of unique surface protein markers on DA-seq clusters and markers that are differentially expressed between RPESC-RPE and PSC-RPE. (e) Flow cytometry analysis showing CD55 expression in RPESC-RPE and PSC-RPE from the CITE-seq dataset (n = 3 each for RPESC-RPE and PSC-RPE). (f) Scatter plot with mean for percentage of cells positive for CD55 in RPESC-RPE (mean = 54.57%) and PSC-RPE (mean = 30.03%) assayed using flow cytometry (p value >0.05). (g) Representative images of 10-week RPESC-RPE and PSC-RPE cells stained with fluorophore-conjugated clone of the CITE-seq antibody against TNFRSF14 (scale bar = 20μm).

With improved separation of RPESC-RPE and PSC-RPE clusters in the multimodal dataset, we next examined the expression of several key genes that involved in retinal and RPE development (**Supplementary Figure 2a**). *PAX6*, a neuroectoderm marker, was highly expressed in PSC-RPE clusters P4, P5, and P7, which also exhibited modest *VSX2* expression, particularly in P5. *SOX2*, a retinal progenitor marker, was also enriched in P5. In contrast, *RAX*, a retina specification gene, was exclusively detected in adult RPESC-RPE clusters A14, A21, and A23. *LHX2* was expressed in both cell types, while *OTX2*, essential for RPE development and function, was broadly expressed in all RPESC-RPE clusters and most PSC-RPE clusters, with the lowest levels in P4, P5, and P7 (**Supplementary Figure 2b**).

To further characterize RPESC-RPE and PSC-RPE clusters, we analyzed the detected surface proteins in DA-seq clusters using the FindAllMarkers function on ADT data (**Table S8, Figure 2d**). PDPN, an established RPE marker,^28^ was detected in all clusters, with the highest expression in A19 and A25. A mucin-type transmembrane protein involved in immune regulation via T-cells and macrophages,^35,36^ PDPN may serve as a general marker of RPE identity and purity. CD9, a tetraspanin,^37^ was also broadly expressed, with higher expression in PSC-RPE clusters. CD9 associates with CD81, which plays a role in RPE phagocytosis,^37^ however, the specific role of CD9 in RPE cells remains unclear. The receptor FAS, which has been implicated in photoreceptor loss through Fas ligand signaling,^38^ was enriched in cluster P4 and A17. Additionally, class I HLA molecules were elevated in expression in clusters P3, A20, and A22, and *NCR1*, a natural killer (NK) cell receptor involved in cytotoxicity,^39^ was enriched in P9 and A24.

Several surface proteins were uniquely detected in either PSC-RPE or RPESC-RPE clusters (**Figure 2d**). In PSC-RPE, NCAM1, a retinal progenitor marker^40^ was enriched in clusters P5 and P7, which also expressed *PAX6* and *VSX2 (***Supplementary Figure 2a**), suggesting a higher proportion of progenitor-like cells not fully committed to the RPE fate. TFRC, associated with proliferative vitreoretinopathy^41^ and iron overload, which has been linked to photoreceptor damage,^42^ was broadly expressed across PSC-RPE clusters. RPESC-RPE clusters also expressed surface markers minimally or not detected in PSC-RPE. CD55, a glycoprotein that accelerates the decay of complement factors to limit cellular damage,^43^ was highly expressed in A21. Given the association between high complement factors and AMD, a molecule that can disrupt the complement cascade may be beneficial, and delivery of CD55 is being explored as a therapeutic for AMD.^43^ CD73 (NT5E) was predominantly detected in several RPESC-RPE clusters, particularly A18. CD73 is an ectoenzyme involved in the generation of adenosine through hydrolysis of adenosine monophosphate.^44^ This conversion has immunosuppressive properties.^45^ These findings highlight source-specific surface markers with potential relevance for cell identity, immune regulation, and therapeutic targeting.

To independently validate surface marker expression, we performed flow cytometry and immunohistochemistry on select markers. CD55, enriched in A21 by CITE-seq, was detectable on both RPESC-RPE and PSC-RPE cells by flow cytometry but its expression was consistently higher across all three adult RPESC-RPE lines, whereas PSC-RPE lines exhibited more variable levels (**Figure 2e,f**), with overall lower expression. We also examined TNFRSF14 (CD270 or HVEM), a TNF receptor superfamily member involved in immune homeostasis via context-dependent modulation of T cell responses^46^ that was enriched in A14 by CITE-seq. Consistent with previous reports of low TNFRSF14 expression in PSC-RPE and fetal RPE,^47^ we observed low levels in PSC-RPE, but notably higher expression in RPESC-RPE using the same fluorophore-conjugated antibody clone used in CITE-seq (**Figure 2g**).

### Enrichment analysis reveals differences in metabolism, immunity, developmental state and stress response

Although overall similarities resulted in mixed transcriptomic clusters of RPESC-RPE and PSC-RPE (Fig 1a), multimodal clustering enabled source-specific separation of the cell types indicating underlying differences in the two cell types. To identify differences between the two cell types, we used the Seurat FindAllMarkers function, grouped by cell source, and identified approximately 6,000 DEGs between PSC-RPE and adult RPESC-RPE (**Table S9**). We then performed GO enrichment analysis using the Single Cell Pathway Analysis (SCPA) tool, which calculates fold changes and q-values for over 3,500 GO-BP terms.^48^ Of these, around 1,500 terms showed statistical significance (adjusted p-value of <0.05), though most had modest fold changes (<2) and qvalues (<3). We focused on GO terms with a fold change >2 and a qval >3 (**Table S10**). Because the qval metric in this analysis pipeline signifies transcriptional change, we used this as an additional guide to identify the top 10 GO categories, representing different functional processes, that were enriched in RPESC-RPE and PSC-RPE (**Figure 3a,b**).

**Figure 3:**
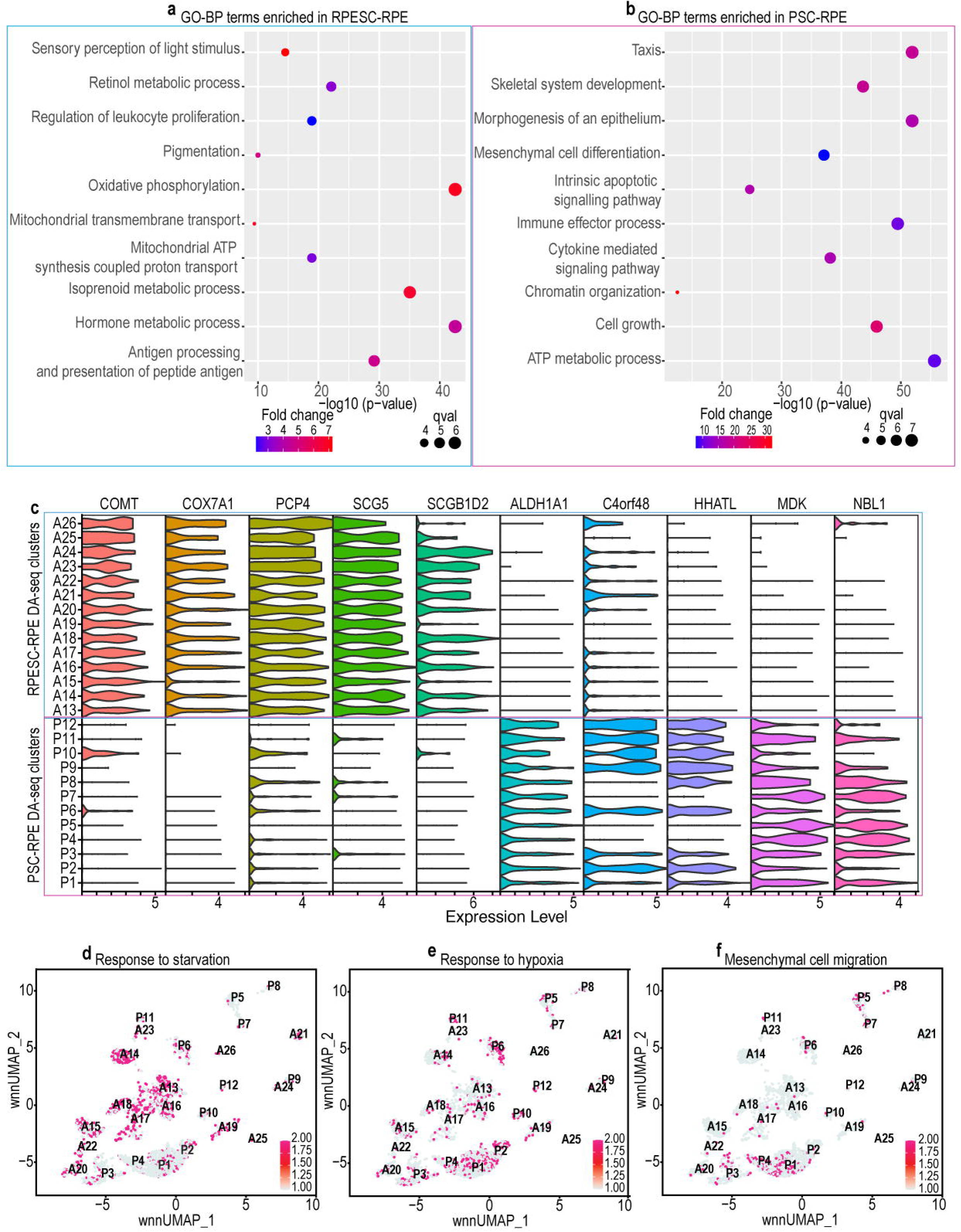
Pathway enrichment and transcript expression of RPESC-RPE and PSC-RPE. (a) Pathways enriched in RPESC-RPE cells related to GO Biological Process terms. (b) Pathways enriched in PSC-RPE cells related to GO Biological Process terms. (c) Stacked ViolinPlot of unique and highly expressed transcripts in RPESC-RPE and PSC-RPE across DA-seq clusters. (d) FeaturePlot showing distribution of cells with enriched genes annotated to GO-BP term “response to starvation” in DA-seq clusters. (e) FeaturePlot showing distribution of cells with enriched genes annotated to GO-BP term “response to hypoxia” in DA-seq clusters. (f) FeaturePlot showing distribution of cells with enriched genes related to GO-BP term “mesenchymal cell migration” in DA-seq clusters. The color scale in panels (d), (e), and (f) represents gene expression for the annotated genes in individual cells based on module scores derived from the AddModuleScore function.

RPESC-RPE cells were enriched for GO terms related to the visual cycle (e.g., *retinol metabolism*, *sensory perception of light stimulus*), mitochondrial function (*oxidative phosphorylation*, *ATP synthesis*), metabolism (e.g., *phenol compound metabolism*, *hormone metabolism, fatty acid metabolic process*), pigmentation, and immune response (**Figure 3a**). Indeed, we observed significantly higher expression of several key RPE genes in adult RPESC-RPE cells, including *MITF, RPE65, SERPINF1, BEST1, ENPP2,* and *LRAT* (**Supplementary Figures 2c-h**). In contrast, PSC-RPE cells showed enrichment for developmental and progenitor-related GO terms, including *chromatin organization*, *epithelial morphogenesis*, *mesenchymal cell differentiation*, and *cell growth*, as well as taxis-related terms (**Figure 3b**). These findings align with a more immature transcriptional state for PSC-RPE than adult RPESC-RPE, and potentially greater migration capacity in PSC-RPE.

Both cell types showed enrichment in gene categories that are related to energy metabolism, however, the specific GO terms differed. RPESC-RPE were enriched for mitochondrial-specific energy production, while PSC-RPE showed broader enrichment in ATP hydrolysis and regulation. This suggests distinct energy kinetics between the two RPE sources (**Figure 3a,b**). Enrichment of immune-related pathways in both PSC-RPE and RPESC-RPE is consistent with the known immunomodulatory role of RPE^49^ (**Figure 3a,b**). However, the specific enrichment terms related to immune function in RPESC-RPE (*regulation of leukocyte proliferation*, *antigen processing and presentation of peptide antigen)* and PSC-RPE (*immune effector process*, *cytokine mediated signaling pathway*) suggest differences in immune responses between the two cell types, which warrants further investigation.

To understand how these differences mapped onto the multimodal clusters, we focused on two key RPE functions that showed differential expression “*fatty acid metabolic process”* and “*sensory perception of light stimulus.”* Using these gene sets, we calculated module scores across DA-seq clusters. Clusters P4, P5, and P7 (PSC-RPE) showed negative scores for both terms, indicating reduced expression. In contrast, clusters P2, A19, and A25 had the highest scores, and enrichment was overall greater in RPESC-RPE than PSC-RPE (**Supplementary Figure 3a,b; Table S11**). These results confirm that multimodal clustering preserves both cellular identity and functional heterogeneity within RPE populations.

Interestingly, this analysis also identified several unique or highly enriched genes for each cell type (**Figure 3c, Table S9**). *COMT*, involved in dopamine metabolism,^50^ *COX7A1*, involved in oxidative phosphorylation,^51^ *PCP4,* a calmodulin-binding protein that acts as a calcium sensor,^52^ *SCG5,* a secreted chaperone that prevents protein aggregation,^53^ and *SCGB1D2*, a secretoglobin,^54^ were strongly detected in RPESC-RPE but not in PSC-RPE. *ALDH1A1*, an aldehyde dehydrogenase,^55^ *C4orf48*, a neuropeptide,^56^ *HHATL*, involved in the post-translational modification of sonic hedgehog,^57^ *MDK*, a growth factor involved in neurogenesis,^58^ and *NBL1*, a BMP antagonist and tumor suppressor,^59^ showed the reverse pattern. It will be worthwhile investigating the function of these enriched genes in each cell type.

A characteristic of AMD is drusen deposits, which are composed of lipoproteins and pigment molecules, between RPE and the Bruch’s membrane, resulting in thickening of the Bruch’s membrane. This affects the choroidal supply of oxygen and nutrients to RPE and photoreceptors.^60^ We calculated module scores for multimodal clusters using genes annotated to terms “*response to starvation*” and “*response to hypoxia”* to examine if some clusters were primed to respond to nutrient and oxygen deprivation. RPESC-RPE clusters (A13–A26) were enriched for genes associated with *response to starvation* (**Figure 3d**), whereas PSC-RPE clusters (P1–P12), particularly P6 and P10, were enriched for *response to hypoxia* genes (**Figure 3e)**, indicating that each RPE cell may have different stress response profiles.

Bruch’s membrane supports RPE cell attachment and thickening of Bruch’s membrane that occurs with age and in disease pathogenesis can impede RPE attachment.^61^ RPE cells that fail to attach to Bruch’s membrane, when exposed to serum factors due to disruption of the blood-retina barrier, can undergo epithelial-to-mesenchymal transition (EMT) and migrate into the vitreous to form epiretinal membranes (ERM).^62,63^ Studies have also reported ERM formation following PSC-RPE transplantation and these may require surgical intervention.^64–66^ To assess EMT potential, we examined the expression of genes associated with the GO-BP term *mesenchymal cell migration*. PSC-RPE clusters P1 and P4 showed marked enrichment, while RPESC-RPE clusters expressed these genes at low levels (**Figure 3f).** These findings are consistent with a previous report that PSC-RPE have greater propensity to undergo EMT than adult RPESC-RPE.^67^

Together, these findings highlight distinct transcriptional signatures between RPESC-RPE and PSC-RPE, particularly in maturation-related, metabolic, immune, plasticity, and stress response pathways. These differences may influence the functional properties of the two sources of RPE cells post-transplantation.

### Differential adhesion and immune modulatory programs in RPESC-RPE and PSC-RPE

Successful RPE transplantation requires effective engraftment and integration into host retinal tissue, with adhesion to Bruch’s membrane being an important determinant of graft survival.^68^ To assess whether adhesion-related features differed between cell types and clusters, we calculated module scores for “*cell–cell adhesion*” and “*cell–matrix adhesion*” using the *AddModuleScore* function. PSC-RPE had higher module scores for *cell-cell adhesion* (PSC-RPE vs. RPESC-RPE; 0.0683 vs. 0.053), while RPESC-RPE had higher module scores for *cell-matrix adhesion* (PSC-RPE vs. RPESC-RPE; 0.077 vs. 0.085) (**Figure 4a**). When analyzed at the subpopulation level, PSC-RPE clusters P6, P9, and P10, along with RPESC-RPE cluster A26, were most enriched for *cell–cell adhesion*. In contrast, RPESC-RPE clusters A25 and A26 showed the strongest enrichment for *cell–matrix adhesion*, whereas PSC-RPE clusters P5 and P7 had the lowest expression of this gene set. These findings suggest that RPESC-RPE and PSC-RPE may utilize distinct adhesion programs, which may influence their engraftment properties and integration efficiency *in vivo*.

**Figure 4:**
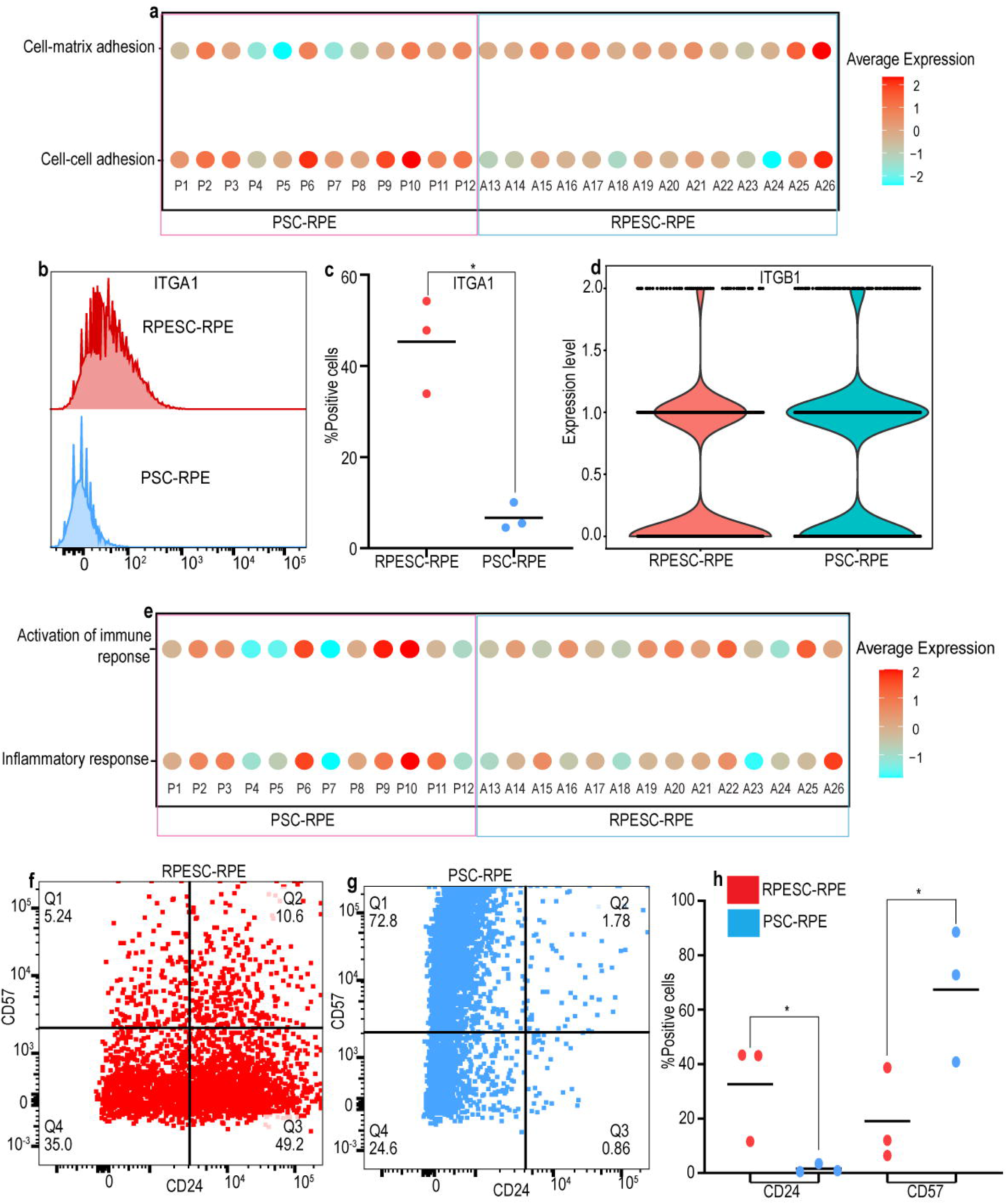
RPESC-RPE and PSC-RPE differ in adhesion and immune response properties. (a) DotPlot showing relative expression of genes related to GO-BP terms “cell-cell adhesion” and “cell-matrix adhesion” in DA-seq clusters based on module scores. (b) Flow cytometry analysis of ITGA1 expression in RPESC-RPE and PSC-RPE from the CITE-seq dataset (n = 3 cell lines each for RPESC-RPE and PSC-RPE). (c) Scatter plot with mean for percentage of cells positive for ITGA1 in RPESC-RPE (mean = 45.30%) and PSC-RPE (6.702%) detected using flow cytometry (*p value <0.05). (d) ViolinPlot showing surface protein expression of ITGB1 in RPESC-RPE and PSC-RPE from the CITE-seq dataset. (e) DotPlot showing relative expression of genes related to GO-BP terms “activation of immune response” and “inflammatory response” in DA-seq clusters based on module scores. Representative flow cytometry data for CD24 and CD57 expression in (f) RPESC-RPE and (g) PSC-RPE. (h) Scatter plot with mean for percentage of cells positive for CD24 and B3GAT1 in RPESC-RPE (CD24: mean = 32.67%; CD57: mean = 19.05%) and PSC-RPE (CD24: mean = 1.623%; CD57: mean = 67.37%) estimated using flow cytometry (n = 3 lines each for RPESC-RPE and PSC-RPE, *p value <0.05).

Given the differences in cell–matrix adhesion molecules, we examined integrin expression using CITE-seq. Interestingly, we found that ITGA1 was uniquely detected on RPESC-RPE, while ITGA2 was detected only on PSC-RPE (**Figure 2d, Supplementary Figure 3c,d**). Flow cytometry confirmed higher ITGA1 protein levels in RPESC-RPE (**Figure 4b,c**). Both integrins pair with ITGB1, which showed no differential expression (**Figure 4d**). Notably, ITGA1 binds preferentially to type IV collagen, the major collagen in Bruch’s membrane,^69^ whereas ITGA2 has a higher affinity for collagen I than IV.^70^ It will be interesting to evaluate whether the expression of ITGA1 enables adult RPESC-RPE to attach more strongly to Bruch’s membrane.

Activation of immune and inflammatory pathways post-transplantation influences graft survival and clinical outcomes.^71^ Our multimodal analysis revealed relatively modest differences in immune-related gene expression between RPESC-RPE and PSC-RPE, with PSC-RPE showing slightly higher module scores for “*activation of immune response*” (PSC-RPE vs. RPESC-RPE; 0.095 vs. 0.092) and “*inflammatory response*” (PSC-RPE vs. RPE; 0.069 vs. 0.063). Subpopulation-level analysis showed that PSC-RPE clusters P9 and P10 were enriched for immune activation genes, while clusters P10 and A24 were enriched for inflammatory response genes. In contrast, clusters P4, P7, and A23 exhibited minimal expression of these gene sets (**Figure 4e**). These subpopulation-level differences suggest that RPE cells may exhibit both immunogenic and immunosuppressive phenotypes after transplantation. Indeed, this has observed for PSC-RPE cells.^72^ Of the immunogenic molecules previously reported on PSC-RPE cells^72^ (e.g., *CXCL9, CXCL10, CXCL11, CCL2, IL6, C3, C5, ICAM1*) and immunosuppressive molecules (e.g., *HLA-E, CD46, CD59, THBS-1 (thrombospondin 1), SERPINF1*), we detected gene expression only for the immunosuppressive markers in both RPESC-RPE and PSC-RPE cells. The immunogenic genes were not detected, which may reflect low expression levels, dependence on specific donor lines or culture conditions, or that these molecules are induced following transplantation.

The immunomodulatory molecules CD24 and CD57 were highly differentially detected between RPESC-RPE and PSC-RPE (**Supplementary Figure 3e,f**). Flow cytometry analysis confirmed this, with up to 45% of RPESC-RPE cells expressing CD24 and about 70% of PSC-RPE cells expressing CD57, across the three lines tested for each cell source (**Figure 4f-h**). CD24 binds to Siglec-10, which can prevent macrophage phagocytosis,^73^ and is part of the “don’t eat me” signaling network that promotes cell survival through immune evasion.^73,74^ CD57 is found on subsets of NK and T cells, where it contributes to suppression of autoimmunity and graft-directed immune responses.^75,76^ Thus, CD24 in RPESC-RPE and CD57 in PSC-RPE may each confer protection against immune rejection, albeit by different mechanisms. Notably, a CD24⁺CD57⁺ double-positive population was observed in adult RPESC-RPE but was much reduced in PSC-RPE (**Figure 4f,g**).

## DISCUSSION

AMD is a progressive, degenerative disorder that severely impacts quality of life.^77^ Limited treatment options have created an urgent need for therapeutic strategies to preserve or restore vision. Cell-based therapies are a promising strategy under active clinical investigation. PSC-derived RPE cells offer a readily renewable, scalable source and are being tested in multiple trials worldwide. Adult RPESC-RPE cells, which are derived from an adult stem cell population that has an inherent specification toward RPE cells as well as a robust expansion capacity, are also in clinical evaluation (Trial ID: NCT04627428). Cell replacement strategies aim to replace lost RPE cells, control disease pathogenesis, and restore vision. The success of cell therapy, therefore, is predicated on the ability of the graft tissue to survive and perform all key RPE functions, as well as mitigate the damage in the eye. While both PSC and adult RPESC sources share common properties such as self-renewal and the ability to give rise to cells with RPE features, comparing the cell products derived from these two cell sources can offer insights into shared features and differences that can aid in identifying predictors of successful clinical outcomes. In this study, we compared transcript and surface protein profiles of cultured RPESC-RPE and PSC-RPE cells using multimodal CITE-seq analysis.

Several studies have compared PSC-RPE with native adult and fetal RPE and identified differences in gene expression, notably finding that PSC-RPE cells are less mature, reaching a developmental stage similar to that of fetal RPE.^28,78,79^ Consistent with these findings, we observed enrichment of developmental pathways in PSC-RPE, along with subpopulation-specific expression of developmental genes and surface proteins. In contrast, adult RPESC-RPE exhibited higher expression of mature RPE markers and genes involved in key functions such as visual pigment recycling. Hence, adult RPESC-RPE cells appear to be more mature overall. Markert et al. proposed that chromatin accessibility networks contribute to PSC-RPE plasticity, and that suppressing these networks may promote their maturation.^67^ It has also been demonstrated that PSC-RPE can attain more mature fates, both with prolonged culture and after transplantation.^22,67,80^ It is possible that the microenvironment encountered by cells engrafting into the retina may affect the functional properties of the transplanted RPE cells. However, given the stressful microenvironment in the AMD eye, initial phenotypic differences may prove important. Monitoring the state of maturation and functionality of the RPE from each source in different models of retinal degeneration and aging will be worthwhile.

RPESC-RPE were enriched for several metabolic terms, including *oxidative phosphorylation, phenol compound metabolism, hormone metabolism* and *fatty acid metabolic process.* The RPE is a highly metabolic tissue, with demands related to constant regeneration of visual pigment, active ion transport, daily phagocytosis and degradation of millions of shed photoreceptor discs, robust mitochondrial respiration, glycolysis, and lipid metabolism.^81–83^ Hence, differences in various metabolic pathways between the source RPE cells may contribute to different functional attributes post-transplantation. For example, RPE and photoreceptors form a metabolic duo, with photoreceptors utilizing glucose and generating lactate as a byproduct, which is then converted to ATP in the RPE through oxidative phosphorylation.^84^ A metabolic shift to increased glycolysis in RPE can result in photoreceptor degeneration,^84^ therefore enhanced *oxidative phosphorylation* may be protective. *Fatty acid metabolism* is tightly coupled between the photoreceptors and the RPE. The products of mitochondrial β-oxidation fuel the TCA cycle and provide adenosine triphosphate (ATP), which supports the demands of RPE cells; again, increases in this pathway may benefit transplant function and survival. *Phenol compound metabolism* is also enriched in RPESC-RPE. Phenol compounds are potent antioxidants and can protect RPE cells from oxidative stress-induced damage.^85–87^

Transcriptomic data revealed a higher EMT and mesenchymal migration signature in PSC-RPE, suggesting an elevated risk of ERM formation, consistent with prior reports.^67^ Following RPE transplantation in the clinic, ERM formation can occur and should be monitored in case surgical removal is necessary.^88,89^ We observed increased expression of *C4orf48,* which codes for an mRNA-binding secreted micropeptide^56^, in PSC-RPE. Although little is known about the role of *C4orf48* in RPE, it has been implicated in TGF-β1–mediated renal fibrosis via the transferrin receptor (TFRC).^56^ It is interesting to note that both *C4orf48* and TFRC expression are particularly enriched in all PSC-RPE clusters, supporting a potential mechanism for TGF-β1-induced ERM formation in PSC-RPE.^67^ While inhibition of the p38-MAPK pathway has been proposed as a mechanism to overcome this risk,^67^ alternative strategies may include TFRC modulation or selective depletion of high-risk RPE clusters based on surface protein expression.

Our data show distinct expression of immune-related surface molecules on RPE cells from each source. Native RPE cells exhibit immunomodulatory properties and express complement regulatory proteins such as CD55 and CD59, which protect against complement-mediated damage.^49^ Due to these properties and the immune-privileged nature of the subretinal space, allogeneic RPE transplantation is considered relatively safe, as supported by clinical trials.^88^ Notably, Kashani et al. demonstrated survival of embryonic stem cell-derived RPE up to two years post-transplantation, despite immune cell infiltration, with the graft retaining RPE marker expression.^90^ Nevertheless, as shown in the rhesus monkey, allogenic PSC-RPE cells injected into the subretinal space can be actively rejected, indicating that understanding the immunomodulatory characteristics of the RPE is valuable to protect graft health over the long-term.^47,72,91,92^ In our study, CD24 and CD57 showed distinct expression patterns on RPESC-RPE and PSC-RPE, respectively, and this may contribute to immune protection post-transplantation. Some PSC-RPE derivation protocols deplete CD24⁺ cells,^93^ which may inadvertently reduce immunosuppressive capacity, given its role as a “don’t eat me” signal.^74^ Conversely, PSC-RPE cells express robust CD57, which can also prevent graft rejection.^76^ RPESC-RPE show an enriched expression of TNFRSF14, which has been shown in other cell types to be pro-inflammatory or immune-suppressing depending on context.^94^ Hence, it will be interesting to examine the role of TNFRSF14 in RPE engraftment. Overall, our data suggest that RPESC-RPE and PSC-RPE cells employ distinct pathways for immune modulation and likely exhibit differences in susceptibility to graft rejection, depending on the subretinal environment. Given that over 1,500 genes are annotated to the GO-BP terms “activation of immune response” and “inflammatory response,” our findings underscore the need for a more comprehensive evaluation of the immune-regulatory genes and proteins expressed by RPE cells and how these are modulated by the subretinal environment encountered in the AMD eye.

In conclusion, our study underscores that there are diverse RPE subtypes within cultured RPESC-RPE and PSC-RPE. While these share critical features, including robust expression of RPE signature genes, they also have notable differences in morphology, gene expression, and surface protein profiles. These distinctions may influence post-transplantation performance. By leveraging CITE-seq to identify surface markers and flow cytometry to isolate specific clusters, we can begin to dissect the functional heterogeneity of RPE. Enriching for clusters that enhance adhesion, immune tolerance, or other aspects of RPE functionality holds promise for improving the success of RPE cell replacement therapies.

### Limitations of the study

One limitation of this study is the use of trypsin to generate single-cell suspensions for CITE-seq, which may alter surface epitopes and therefore affect the comparative analyses. Additionally, the current antibody panel is limited, with many targets derived from hematopoietic lineages. Our findings underscore the importance of expanding CITE-seq panels to encompass a more comprehensive range of RPE-relevant surface markers.

## ACKNOWLEDGEMENTS

We are grateful to the Regenerative Research Foundation (RRF) for support, the NSCI NeuraCell core facility for iPSC-RPE production and technical support.

**hPSC lines** (**Table S1**) were used in this study. The authors acknowledge the invaluable contributions of the tissue donors and their families and the assistance of the support staff to generate these lines. B82.1, B88.2 and R255.1 pluripotent stem cell lines^95^ and differentiated PSC-RPE are available through Neuracell (https://www.neuracell.org/ or by contacting). The CRISPRi-ready clone of the WTC11 line was a kind gift from Martin Kampmann (UCSF).^96^

## Author contributions

Conceptualization, NCB, ST, BLA, JHS; Methodology, FF, NCB, ST; Software, FF, SS, NCB; Formal Analysis, SN, FF, NCB, ST, BLA; Investigation, SN, FF, TB, SL, YW, JAK; Resources, ST, JHS; Data Curation, SN, NCB, ST, BLA; Writing - Original Draft, SN, ST, BLA, NCB; Writing – Review and Editing, NCB, BLA, ST; Supervision, NCB, ST, JHS; Project Administration, NB, BLA, ST; Funding Acquisition, ST, NCB, BLA, JHS.

## Lead contact

Request for further information and resources should be directed to and will be fulfilled by lead contact Sally Temple.

## Material availability

Human PSC lines and PSC-derived RPE will be made available on request and upon completion of a materials transfer agreement if there is potential for commercial application.

## Data and code availability

Single-cell RNA seq data generated during this study are available at GEO: GSE244572. All code utilized for analysis can be found at: https://anonymous.4open.science/r/Adult_vs_Pluripotent_RPE-C622/

## METHODS

### Eye dissection and RPESC-RPE cultures

Human adult RPE cells were isolated from human donor eyes and maintained in culture as reported previously.^21,97^ All donor eyes were obtained from approved eye banks after receiving consent for research use. Donor details are listed in Table S1. Donor globes were immersed in betadine solution for 5 minutes and washed with 1X PBS three times. The RPE layer was exposed by making a cut at the ora serrata and removing the anterior portion, vitreous humor, and the retina. The posterior cups were incubated with collagenase IV (1mg/ml) at 37°C incubator and dissociated RPE cells were collected and cultured on synthemax (Corning)-coated plates in RPE medium containing DMEM F12 50/50 medium (Corning), MEM alpha modification medium (Sigma-Aldrich), 1.25 ml Glutamax (Gibco), 2.5 ml sodium pyruvate (Gibco), 2.5 ml niacinamide (1 M; Spectrum Chemical Inc.), 2.5 ml MEM non-essential amino acid solution (Gibco), 10% heat-inactivated fetal bovine serum (FBS), supplemented with THT (taurine, hydrocortisone, triiodo-thyronin), and 1.25 ml N1 or N2 medium supplement (Sigma-Aldrich) a humidified incubator at 37°C and 5% CO2. The cells were shifted to 2% FBS containing RPE media after the first week of plating and medium was replaced three times a week for the entire duration of culture (passage P0). After 7 weeks, the cells were passaged onto synthemax-coated plates and cultured for an additional 7-8 weeks (passage P1). P1 RPESC-RPE cells were cryopreserved in CryoStor CS2 (Stem Cell Technologies). For experiments in this study, RPESC-RPE cryopreserved cells were thawed and plated at 100,000 cells/well on a synthemax-coated 24-well plate in 10% FBS containing RPE media. Cells were maintained in 2% FBS containing RPE media after the first week and were cultured for 10 weeks in a humidified incubator at 37°C and 5% CO2. The medium was replaced three times a week.

### PSC-RPE Differentiation and Culture

PSC-RPE cells derived from three different hPSC lines using a modified version of a previously published differentiation protocol^24^. Details about the hPSC lines are listed in Table S1. The hPSC lines were maintained as colonies on cultrex-coated dishes until they were 60-70% confluent and then harvested as single cells using TrypLE express with Phenol Red. The cells were counted and plated at 300,000 cell/cm^2^ of a cultrex-coated 24-well plate with ROCK inhibitor. Cells were allowed to attach overnight, and if the cells were 100% confluent, differentiation media was introduced and was counted as day 0 of differentiation. Base media for differentiation consisted of DMEM F12 (ThermoFisher Scientific) with 2.5 ml MEM non-essential amino acids, 1X N2 supplement (ThermoFisher Scientific) and 1X B27 supplement without vitamin A (ThermoFisher Scientific). Cells received base media supplemented with LDN193189, DKK1, IGF1, and niacinamide for the first four days, with FGF2 being introduced on day 2. On days 4 and 5, cells were supplemented with DKK1, IGF1 and activin A in base media. From days 6 onwards, cells were maintained in base media containing activin A and SU5402, with the introduction of CHIR on day 8. The combination of activin A, SU5402, and CHIR was continued until day 18 of differentiation. Media was replaced every day for the duration of differentiation. From day 19 onwards, cells were maintained in the same RPE media used to culture RPESC-RPE (passage P0). Cells were passaged between days 40-45 on to synthemax-coated dishes and were maintained in RPE media until days 65-70 (passage P1), after which they cryopreserved in CryoStor CS2. For experiments in this study, PSC-RPE cryopreserved cells were thawed and plated at 100,000 cells/well on a synthemax-coated 24-well plate in 10% FBS containing RPE media. Cells were maintained in 2% FBS containing RPE media after the first week and were cultured for 10 weeks in a humidified incubator at 37°C and 5% CO2. The medium was replaced three times a week.

### CITE-Seq and library preparation for scRNA-seq

Samples were prepared for scRNA sequencing using the ICELL8 system as described previously.^97^ RPE cells from the RPESC and PSC sources were dissociated using 0.25% trypsin to obtain a single cell suspension and then stained with the oligo-conjugated antibodies. These single cells were stained with SYTO64 and dispensed onto iCELL8 3’ DE chips using the multisample NaNoDispenser (Takara Bio). Wells containing doublet cells, based on SYTO64 staining, were not used for cDNA and library preparation. Reverse transcription PCR was performed with lysed cells in the chips using the 3’ DE reagent kit (Takara Bio) per manufacturer’s instructions. For ADT sequences, an ADT additive primer (0.1 ng) was included during RT-PCR. cDNA was collected, concentrated, and purified using AMPure XP magnetic beads (Beckman Coulter) per manufacturer’s instructions. Nextera XT DNA library preparation kit (Illumina) was used for cDNA library generation based on instructions from Takara’s 3’ DE chip and reagent kit (Biolegend). ADT libraries were amplified using RPI-X primers containing the P7 sequence. Both the cDNA and ADT products were assessed for quality using an Agilent High Sensitivity DNA Kit and Agilent 2100 Bioanalyzer (Agilent Technologies, Palo Alto, CA).The libraries were sequenced on a NovaSeq 6000 high-output flow cell generating 2 x 150bp read lengths.

### Data processing and analysis

Raw Illumina read BCL Files were converted to fastq files using the bcl2fastq2 software (bcl2Fastq v2.19.1, Illumina, Inc). Fastq files were then merged into read1 (ICELL8 barcode sequence) and read2 (transcript sequence) fastq files. Reads were mapped to human hg38 genome using the Cogent NGS Analysis Pipeline (V1.5, Takara Bio) utilizing the STAR aligner.^98^ The resulting transcript and ADT read count matrices were used as the input for further analysis with Seurat (V4 and V5) package for R normalization was performed using the SCTransform pipeline.^99^ To analyze the antibody-derived tag (ADT) data, we devised a novel algorithm to normalize and transform the data utilizing an arcsinh transformation.^100^ We then analyzed the normalized and transformed ADT and raw transcriptomic data using the Seurat package. The RNA counts matrix was converted to a Seurat object using CreateSeuratObject function with a minimum cell cutoff of 3 and a minimum feature cutoff of 200. Data integration and Weighted Nearest Neighbor Analysis were performed using the Seurat package to combine different datasets and cluster cells using both transcriptomic and surfaceome data. All code utilized for analysis can be found at: https://anonymous.4open.science/r/Adult_vs_Pluripotent_RPE-C622/

### Flow Cytometry

Flow cytometry analysis for surface markers was carried out on live cells for both RPESC-RPE and PSC-RPE. Cells were harvested using 0.25% trypsin-EDTA with DNAseI and resuspended in 5% FBS containing PBS to neutralize the trypsin and pelleted down at 300g for 5 minutes. Cells were stained with fluorophore conjugated primary antibodies at a 1:100 dilution in 2.5% FBS containing PBS for 25-30 minutes at room temperature in the dark (APC anti-human CD49a, catalog# 328313, BioLegend; APC anti-human CD55, catalog# 311311, BioLegend; Pe-Cy7 anti-human CD24, catalog# 31119, BioLegend; APC anti-human CD57, catalog# 393305, BioLegend). For experiments with anti-CD24 and anti-CD57, both RPESC-RPE and PSC-RPE were stained simultaneously with the two antibodies at a 1:100 dilution for each antibody. The cells were washed twice and pelleted down at 300g for 5 minutes. Cells were resuspended in 300ul 2.5% FBS containing PBS for flow cytometry analysis. Staining for CD59 was incorporated as a control because of its ubiquitous expression on RPE cells.^101,102^ Positive gating utilizing anti-CD59 antibodies (APC anti-human CD59, catalog# 304711, BioLegend; Pe-Vio770 anti-human CD59, catalog# 130-129-882, Miltenyi Biotech; BD Horizon BV421anti-human CD59, Fisher Scientific, catalog# 565982) was used to detect cell populations of interest.

### Immunostaining

Cultured cells on 24-well or 96-well plates were fixed for 20 minutes with 4% paraformaldehyde at room temperature. The cells were then washed with Calcium/Magnesium free-DPBS three times. Cells were blocked in 3% BSA for 30 minutes at room temperature. Fluorophore conjugated primary antibodies (PE anti-human CD270, catalog# 318805, BioLegend) diluted to 1:20 in 1X DPBS were used for staining the cells overnight at 4°C in the dark. The following day the cells were washed three times in DPBS. Cells were counterstained with DAPI diluted 1:1000 in DPBS and imaged on Zeiss Axio observer D1 microscope.

### Image analysis

Morphological analysis was performed by analyzing phalloidin-stained images from three different RPESC-RPE and PSC-RPE lines cultured for 10 weeks. Images were converted to grayscale and cell–cell junctions were detected using the Ridge Detection plugin for ImageJ (https://ieeexplore.ieee.org/abstract/document/659930) based on phalloidin staining. Object identification and morphological analysis were performed using the EBImage package^103^ in R.

### Statistical Analyses

Single cell analysis was performed using Seurat V4 and V5 and the Wilcoxon rank sum test was applied to identify differentially expressed genes in all cases where the Seurat FindAllMarkers function was used. For pathway enrichment analysis using ClusterProfiler, false discovery rate was used to account for multiple comparisons. For pathway enrichment analysis using single cell pathway analysis (SCPA), default parameters for p-value correction and multivariate analysis were applied. Student’s t-test was used for analysis of flow cytometry data for RPESC-RPE and PSC-RPE comparisons.

---

**Supplementary Figure 1:**
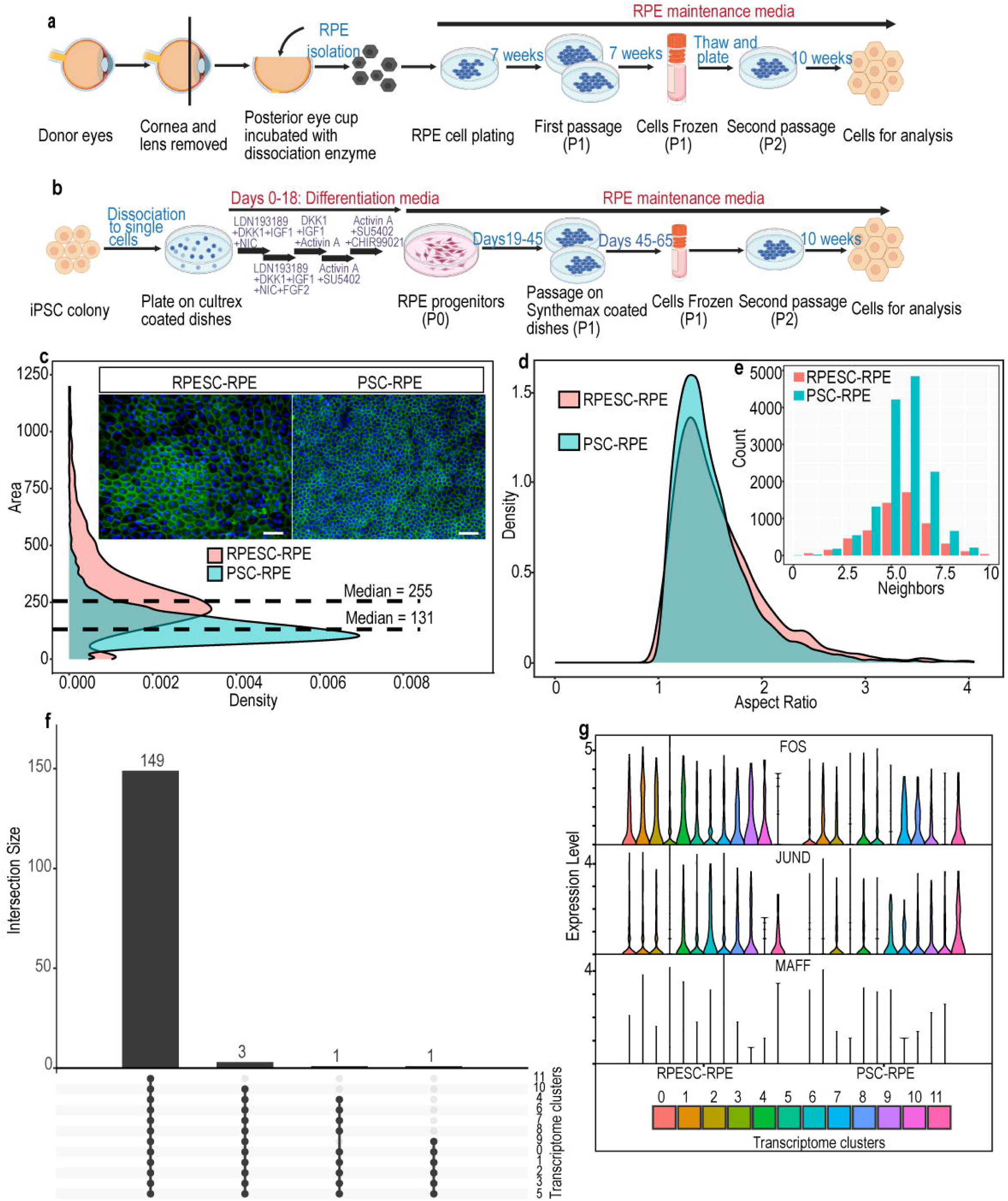
Culture conditions and morphological attributes of RPESC-RPE and PSC-RPE. Schematic of (a) RPESC-RPE culture method. (b) PSC-RPE differentiation strategy. (c) The density plot for surface area in pixels of RPESC-RPE and PSC-RPE. Representative images of F-actin-stained PSC-RPE and RPESC-RPE (F-actin in green and DAPI in blue, scale bar = 100μm) demonstrating differences in cell sizes between the two cell types (n = 3 lines each for RPESC-RPE and PSC-RPE). (d) Density plot comparing aspect ratio (AR) between RPESC-RPE and PSC-RPE. AR indicates cell regularity, varying from 1 (highly regular, expected from mature hexagonal RPE cells) to 3 (highly elongated cells) (n = 3 lines each for RPESC-RPE and PSC-RPE. (e) Grouped Bar Plot showing number of neighbors for RPESC-RPE and PSC-RPE. Hexagonal cobblestone RPE cells are expected to have around 5-7 neighbors (n = 3 lines each for RPESC-RPE and PSC-RPE). (f) Upset plot showing overlap in RPE signature genes across single clusters based on transcriptomics. (g) Stacked ViolinPlot showing expression of transcription factors reported to be expressed in mature RPE cells.

**Supplementary Figure 2:**
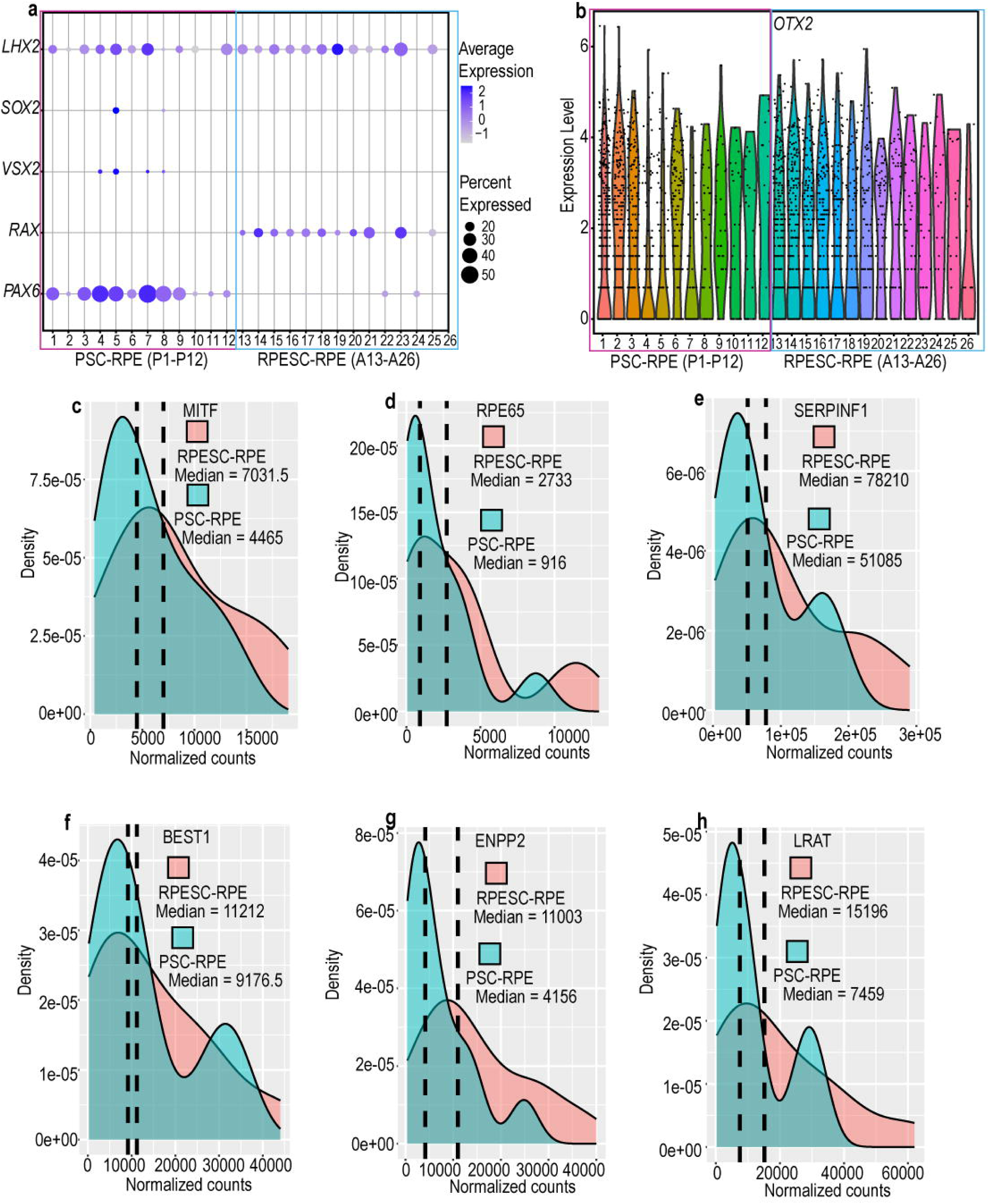
RPESC-RPE and PSC-RPE show differences in expression of genes involved in RPE specification and development. (a) DotPlot showing expression of markers involved in the determination and differentiation of retinal cell fates in DA-seq clusters of RPESC-RPE and PSC-RPE. (b) ViolinPlot showing expressing of *OTX2* across DA-seq clusters of RPESC-RPE and PSC-RPE. Density Plot showing differences in expression of RPE markers grouped by cell source (c) *MITF* (d) *RPE65* (e) *SERPINF1* (f) *BEST1* (g) *ENPP2* (h) *LRAT* in RPESC-RPE and PSC-RPE.

**Supplementary Figure 3:**
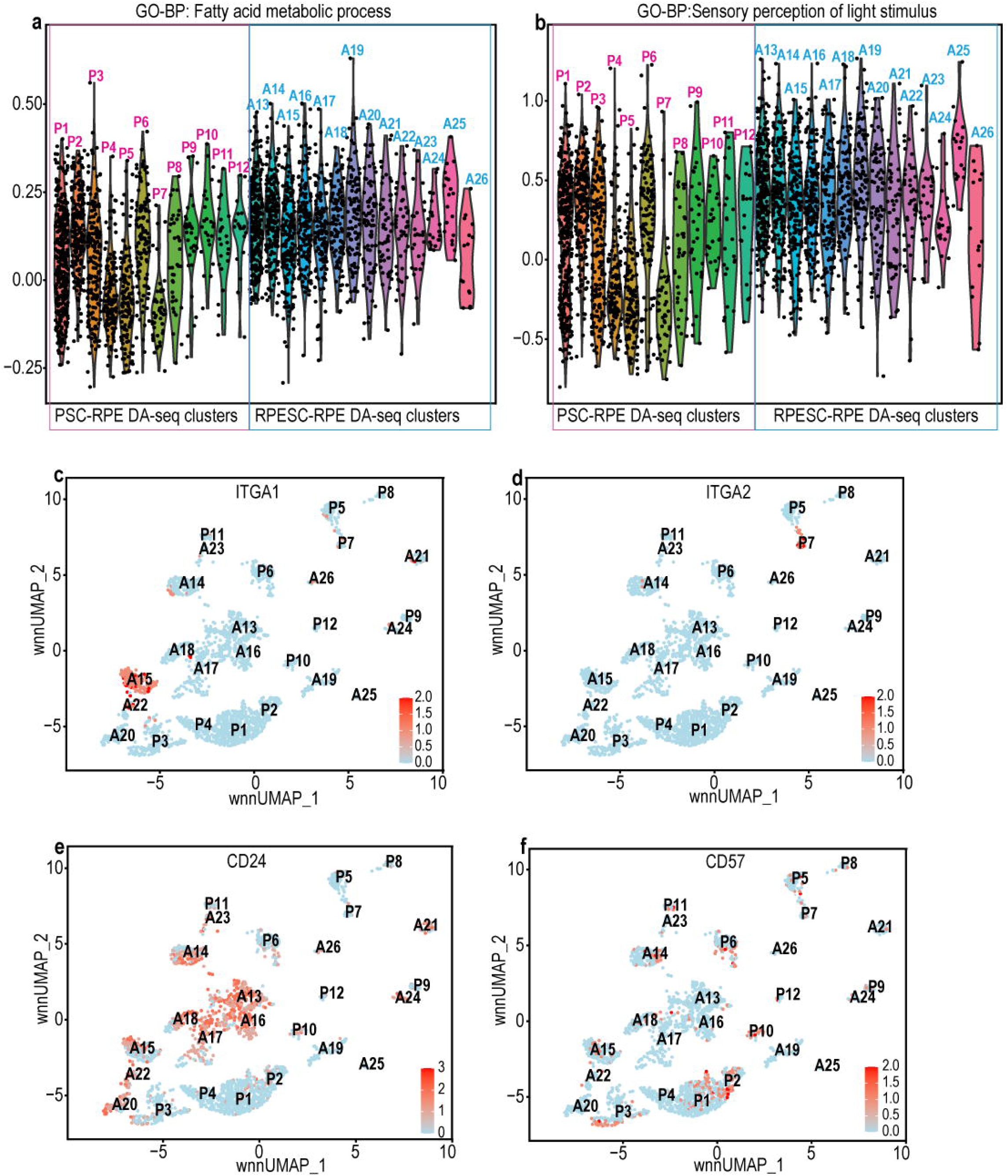
Gene and protein expression of select markers across DA-seq clusters of RPESC-RPE and PSC-RPE. (a) ViolinPlot showing relative expression of genes annotated to GO-BP term “fatty acid metabolic process” across DA-seq clusters based on module scores. (b) ViolinPlot showing relative expression of genes related to GO-BP term “sensory perception of light stimulus” across DA-seq clusters based on module scores. (c) FeaturePlot showing surface protein expression of ITGA1 in DA-seq clusters (color scale = ITGA1 expression in individual cells) (d) FeaturePlot showing surface protein expression of ITGA2 in DA-seq clusters (color scale = ITGA2 expression in individual cells). (e) FeaturePlot showing surface protein expression of CD24 in DA-seq clusters (color scale = CD24 expression in individual cells). (d) FeaturePlot showing surface protein expression of CD57 in DA-seq clusters (color scale = CD57 expression in individual cells).

## Supplementary Tables

**Table S1**: Donor and cell line information for RPESC-RPE and PSC-RPE

**Table S2**: Morphology analysis of RPESC-RPE and PSC-RPE cells

**Table S3**: List of CITE-seq antibodies

**Table S4**: Transcript counts for macular and peripheral RPE markers across transcriptome-based clusters (0-11)

**Table S5**: RPE signature genes by transcriptome-based clusters (0-11)

**Table S6**: List of differentially expressed genes for transcriptome-based clusters (0-11)

**Table S7**: List of enriched pathways across transcriptome-based clusters (0-11) identified using R package ClusterProfiler

**Table S8:** List of signature proteins identified on DA-seq clusters based on multimodal clustering of RPESC-RPE and PSC-RPE

**Table S9**: List of signature genes in RPESC-RPE and PSC-RPE following multimodal clustering

**Table S10**: SCPA output for differentially enriched pathways across RPESC-RPE and PSC-RPE clusters following multimodal clustering

**Table S11**: Mean module scores for each DA-seq cluster calculated using AddModuleScore utilizing gene sets for GO-BP terms “fatty acid metabolic process” and “sensory perception of light stimulus”

